# Spectro-temporal neural dynamics during sentence completion

**DOI:** 10.1101/2021.07.01.450734

**Authors:** Tim Coolen, Alexandru Mihai Dumitrescu, Vincent Wens, Mathieu Bourguignon, Gustavo Lucena Gómez, Antonin Rovai, Niloufar Sadeghi, Charline Urbain, Serge Goldman, Xavier De Tiège

**Affiliations:** Laboratoire de Cartographie fonctionnelle du Cerveau, ULB Neuroscience Institute (UNI), Université libre de Bruxelles (ULB), Brussels, Belgium; Department of Radiology, CUB - Hôpital Erasme, Université libre de Bruxelles (ULB), Brussels, Belgium; Magnetoencenphalography unit, Clinics of Functional Neuroimaging, Service of Nuclear Medicine, CUB - Hôpital Erasme, Université libre de Bruxelles (ULB), Brussels, Belgium; BCBL – Basque Center on Cognition, Brain and Language, 20009 San Sebastian, Spain; Laboratory of Neurophysiology and Movement Biomechanics, ULB Neuroscience Institute (UNI), Université libre de Bruxelles (ULB), Brussels, Belgium; UR2NF, Neuropsychology and Functional Neuroimaging Research Unit, Centre for Research in Cognition and Neurosciences (CRCN), ULB Neuroscience Institute (UNI), Université Libre de Bruxelles(ULB), Brussels, Belgium

**Keywords:** Sentence completion, Semantic Network, Ventral Stream, Magnetoencephalography, Oscillations

## Abstract

This magnetoencephalography (MEG) study aimed at characterizing the spectro-temporal dynamics of brain oscillatory activity elicited by sentence completion (SC). For that purpose, we adapted a version of the SC experimental paradigm typically used in functional magnetic resonance imaging to MEG investigation constraints. Twenty right-handed healthy young adults underwent MEG recordings while they were sequentially presented with short sentences divided in three parts: the first two giving context and the last requiring completion. MEG data were then analysed using a prior-free, non-parametric statistical approach with stringent control of the family-wise error rate. We identified three successive significant neural response patterns associated with distinct spatial and spectro-temporal characteristics: (i) an early (<300 ms) bioccipital 4-10-Hz event-related synchronization (ERS); (ii) an intermediate (at about 400 ms) 8-30-Hz event-related desynchronization (ERD) in an extended semantic network involving the ventral language stream as well as bilateral posterior nodes of the default mode network (DMN) in both hemispheres; (iii) a late (>800 ms) 8-30 Hz ERD involving the left dorsal language stream. Furthermore, the left component of the ventral language stream displayed prolonged ERD after 800 ms compared to the right which showed signs of inhibition in the form of ERS. Overall, this study elucidates the dynamics of the recruitment of the language network that accompany SC and the spectro-temporal signature of an extended semantic network. This MEG adaptation of an SC paradigm also paves the way for novel approaches in presurgical language mapping and may help to understand the neural underpinnings of the alterations of sentence completion in various neurologic disorders affecting language.

## 1. Introduction

Sentence completion (SC) is a well-documented language task in the functional magnetic resonance imaging (fMRI) literature (e.g., Ashtari et al., 2005; Barnett et al., 2014; Black et al., 2017; Kircher et al., 2001; Petrella et al., 2006; Salek et al., 2017; Wilson et al., 2017; Zaca et al., 2012; Zacà et al., 2013). It is endorsed by the American Society of Functional Neuroradiology as a first-choice experimental paradigm for language functional brain mapping (Black et al., 2017) that is typically constructed as a block design (for a description of fMRI designs, see, e.g, Amaro & Barker, 2006). In such design, the active condition usually consists of short written sentences with a missing final part that participants are asked to complete, while the control condition includes low-level visual stimuli such as gibberish written sentences with a missing final part. Likely due to the high linguistic complexity, inherent to sentence-level processing (Vigneau et al., 2006), SC robustly (Black et al., 2017; Zacà et al., 2013) and reliably elicits increases in blood oxygen level dependent (BOLD) signal in the left perisylvian regions (Wilson et al., 2017) deemed essential to language function (for a review, see, e.g., Tremblay & Dick, 2016).

Owing to its exquisite temporal resolution, magnetoencephalography (MEG) provides the opportunity to investigate the spectral and temporal oscillatory neural dynamics underlying sentence processing within left-lateralized language-related regions. Previous MEG studies (e.g., Halgren et al., 2002; Hultén et al., 2019; Kielar et al., 2015; Lam et al., 2016; Meltzer et al., 2017; Meltzer & Braun, 2011; Piai et al., 2015; Wang et al., 2018) did not investigate the neural oscillatory dynamics that accompany the completion of sentences during SC. Instead, they focused on the semantic integration of incoming words in the sentence context (Halgren et al., 2002; Hultén et al., 2019; Lam et al., 2016; Wang et al., 2018). Other studies investigated the effect of the modulation of the phonological and semantic aspects of the sentence (Meltzer et al., 2017) or of specific linguistic abnormalities such as semantic violations (Kielar et al., 2015). Moreover, the sentence endings were given in those studies, or cued by a picture in one of them (Piai et al., 2015) which focused on production aspects.

This MEG study was therefore performed to characterize the spectral, temporal and spatial dynamics of the neural events associated with the whole SC process. For that purpose, we developed a version of the classic fMRI SC experimental paradigm adapted to the constraints of MEG investigations (for a detailed description, see, e.g., Gross et al., 2013). We first analyzed the whole-brain MEG data acquired in right-handed healthy young adults using a prior-free, non-parametric statistical approach with stringent control of the family-wise error rate. Secondly, we restricted the investigation to the bilateral ventral stream of language (Hickok & Poeppel, 2007) to shed light on the relative contribution of its left and right components to sentence processing. Indeed, in addition to the typically left-sided changes elicited by sentence stimuli in left-sided language-related areas (Tzourio-Mazoyer et al., 2017), both fMRI (e.g., Barnett et al., 2014; Wilson et al., 2017; Zaca et al., 2012) and MEG (e.g., Halgren et al., 2002; Hultén et al., 2019; Kielar et al., 2015; Lam et al., 2016; Meltzer et al., 2017) studies also reported contralateral homologous right-sided activity modulations, notably in right posterior temporal regions (e.g., Kircher et al., 2001; Meltzer et al., 2017). Still, the functional role of these right-sided temporal changes during sentence processing remains poorly understood. They may correspond to context processing (Vigneau et al., 2010) or prolonged maintenance of multiple semantic representations necessary to understand the gist of the sentence (Kircher et al., 2001). We expected that, due its exquisite temporal resolution, MEG will yield novel insights into the different functional roles of left and right components of the ventral language stream through potentially divergent spectro-temporal fingerprints.

## 2. Materials and methods

### 2.1. Subjects

Twenty right-handed healthy adult subjects (mean age: 31.2 ± 8.1 years, range: 22.0 – 51.4 years; 11 females) were included in this study. None of them had a prior history of neurologic, psychiatric or learning disorder, nor MRI contraindication. All subjects were right-handed (93.3 ± 8.8 %; range: 77.8 – 100 %) according to the Edinburgh Handedness Inventory scale (Oldfield, 1971).

All subjects contributed to the study after giving written informed consent. They were given a small financial incentive for their participation. The study received prior approval by the Ethics Committee of the CUB – Hôpital Erasme (Université libre de Bruxelles, Brussels, Belgium; REF: P2017/272)

### 2.2. SC paradigm

A total of 72 simple short sentences in French were created (listed in Supplementary Table 1). Each sentence was divided into three parts. The first part (P1) contained the subject, composed of a common noun and its determiner (mean of 2.04 ± 0.31 words; 10.47 ± 2.59 characters, including spaces; e.g., “The hen”). The predicate was partly stated in the second part (P2; 2.53 ± 0.73 words; 12.36 ± 3.29 characters; e.g., “lays an”) and had to be completed by the participants in the third part (P3), which prompted the completion by a visual clue (“_________.”).

A schematic representation of the paradigm is provided in figure 1. A block structure, typical for a clinical fMRI SC paradigm, served as the canvas for this MEG adaptation and consisted of six “*task*” blocks of 60 s each, alternating with “*rest*” periods of 30 s. Total paradigm duration was 9.6 min. The task blocks of interest contained six sentences. A fixation cross lasting 1 s preceded every sentence, and each part thereof (i.e., P1, P2, P3) was shown for 3 s. The thirty-six sentences were different and randomly picked among the available sentence database for each participant. Rest periods consisted of three sequences that were created to visually resemble the target sentences and also divided in a first and a second part, composed only of “*” characters and spaces, followed by the same empty third part (“_________.”). For added clarity, a 1 s cue followed by a 2 s fixation cross announced the beginning of each block.

**Figure 1:**
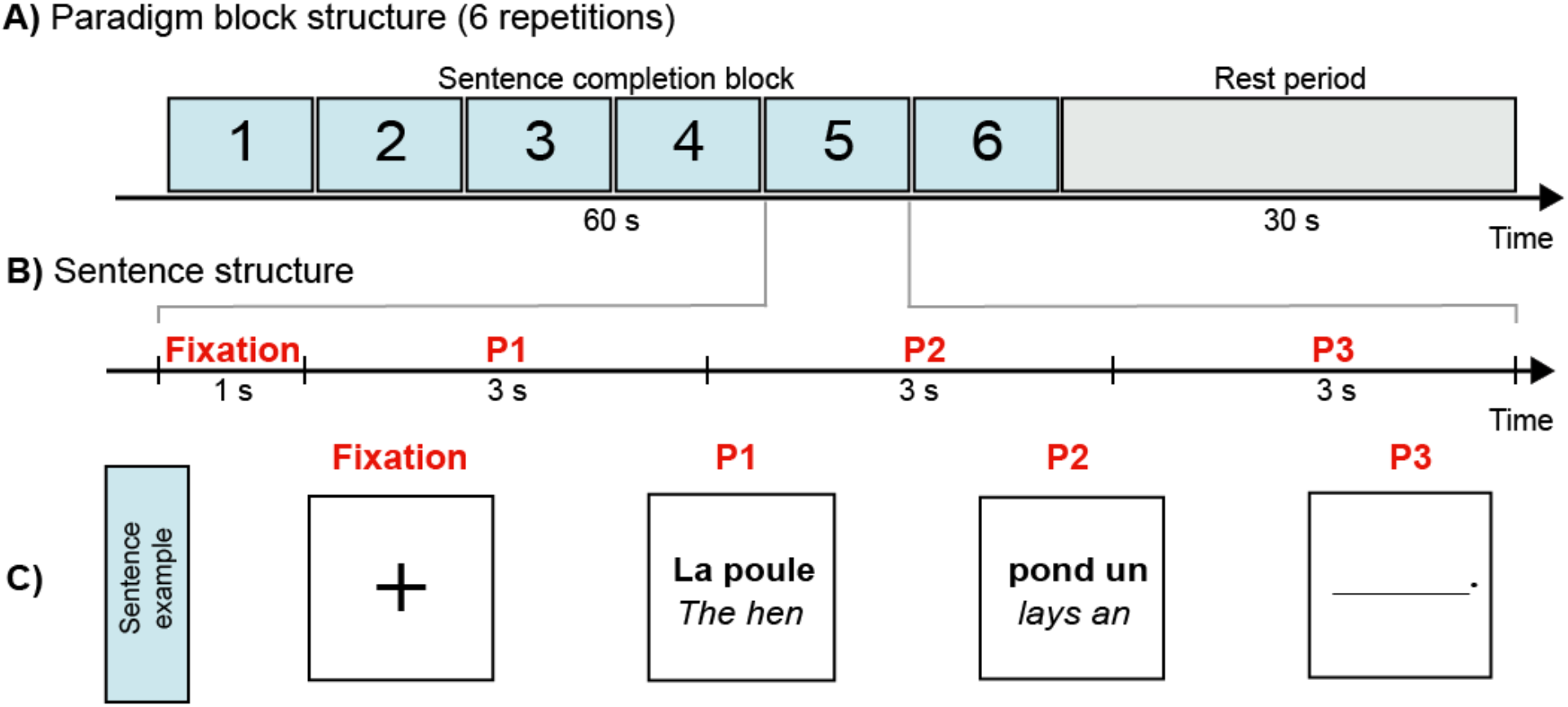
Schematic representation of the SC paradigm. **A)** The general block structure consisted of task blocks of six sentences (blue), alternating with rest periods (grey). **B)** The sentence structure was composed of a fixation cross followed by a first (P1: determiner and noun), second (P2: part of predicate) and third (P3: to complete) part. **C)** Example sentence. The text in French actually presented to the subjects is written in bold. For convenience, the English translation is also provided here in italic.

Subjects were instructed to silently read the first two parts of the sentences and to silently generate the sentence endings with one or few words when presented with the P3 completion cue. During the rest periods, they were asked to simply look at the screen. Prior to the MEG recording, participants were trained on a separate set of six sentences and asked to overtly read and complete the sentences to ensure they correctly understood the instructions. All subjects understood the task and performed the training correctly.

Visual stimuli appeared in white Times New Roman font on a black background. They were projected onto a screen placed at the feet of the MEG bed and made visible to the subject through an oblique mirror so that the visual angle did not exceed 7°. The appearance of each sentence part (P1, P2 and P3) was marked by a specific trigger signal recorded in a separate channel synchronously to the MEG data.

### 2.3. Data acquisition

MEG data were recorded (band-pass: 0.1–330 Hz; sampling rate: 1 kHz) in a lightweight magnetically shielded room (Maxshield™, MEGIN, Croton Healthcare, Helsinki, Finland; see De Tiège et al., 2008 for more details) using a 306-channel whole-scalp-covering neuromagnetometer (Triux™, MEGIN, Croton Healthcare, Helsinki, Finland). Subjects were installed in the supine position to minimize head movement artifacts.

Four fronto-mastoid head-tracking coils monitored the subjects’ head position inside the MEG helmet. The locations of the coils and at least 350 head-surface points (on scalp, nose and face) with respect to anatomical fiducials were recorded with an electromagnetic tracker (Fastrak, Polhemus, Colchester, VT, USA) before starting the MEG session.

For MEG source localization, an anatomical 3D T1-weighted gradient echo sequence (time of repetition = 8.2 ms, time of echo = 3.1 ms, flip angle = 12°, field of view = 24 cm, matrix = 240 x 240, isotropic 1 mm^3^ voxels) of the head was also acquired for each subject using a hybrid 3T SIGNA™ PET-MR scanner (GE Healthcare, Milwaukee, Wisconsin, USA) with a 24-channel head and neck coil.

### 2.4. MEG data preprocessing

Raw MEG data were preprocessed off-line using signal space separation (Taulu et al., 2005) to reduce external interferences and correct for head movements (MaxFilter™ v2.2 with default parameters, MEGIN, Croton Healthcare, Helsinki, Finland). Cardiac, eye-movement and electronic artifacts were then identified by independent component analysis (FastICA algorithm with rank reduction to 30 and nonlinearity *tanh;* Hyvärinen & Oja, 2000; RRID: SCR_013110) applied to sensor time series filtered between 0.5 and 45 Hz (Vigario et al., 2000) and visual inspection of the components (mean number of artifactual components per subject : 2.30 ± 0.57). Artifactual components were then regressed out from the full-rank data.

The preprocessed MEG data were further handled with Fieldtrip (Oostenveld et al., 2011; RRID:SCR_004849). The continuous data were first segmented in 6 s-long epochs covering each sentence part P1, P2 or P3 (2000 ms prestimulus to 4000 ms post-stimulus, each stimulus time being identified by a trigger event). Epochs with absolute signal values exceeding 3 pT in at least one magnetometer or 0.7 pT/cm in at least one gradiometer were discarded as likely contaminated by artifacts.

The time-frequency Fourier coefficients were then estimated using a 7-cycle Morlet wavelet decomposition, for each of the 306 MEG channels (102 magnetometers and 204 planar gradiometers). This was done in a window of interest starting at 500 ms before and ending at 3000 ms after each sentence part (P1, P2, P3) with a 50 ms time increment. Of note, the usage of wider (6 s-long) epochs was necessary to probe the lowest frequencies in this window of interest. The Fourier coefficients were estimated from 1 Hz to 45 Hz by steps of 1 Hz.

### 2.5. Source reconstruction

Individual anatomical MRIs were segmented using the Freesurfer software (Fischl, 2012; Martinos Center for Biomedical Imaging, Massachusetts, USA; RRID: SCR_001847). MEG and MRI coordinate systems were co-registered using the three anatomical fiducial points for initial estimation and the head-surface points to manually refine the surface co-registration.

Individual MEG forward models were then computed using the single-layer Boundary Element Method implemented in the MNE-C software suite (Gramfort et al., 2014; Martinos Center for Biomedical Imaging, Massachusetts, USA; RRID: SCR_005972). To ease the group-level analysis, forward models were based on a source grid obtained from a common 5-mm cubic grid (containing 16102 source locations), built in the Montreal Neurological Institute (MNI) template brain by applying the non-linear spatial deformation estimated with the algorithm implemented in Statistical Parametric Mapping (SPM12, Wellcome Department of Cognitive Neurology, London, UK; RRID: SCR_007037). Three orthogonal current dipoles were placed at each grid point.

The resulting forward models were then inverted via Minimum Norm Estimation (MNE; Dale & Sereno, 1993). The sensor-space noise covariance was estimated from 5 minutes of artifact-free data recorded from an empty room preprocessed using signal space separation and filtered between 0.5 and 45 Hz. The regularization parameter was derived for each condition (P1, P2 and P3) from the signal-to-noise level estimated from the noise covariance and the covariance of the concatenated epoch data (Wens et al., 2015). The depth bias was corrected by noise standardization (Pascual-Marqui, 2002). The MNE inverse operator was then used to project the sensor-level time-frequency Fourier coefficients at each epoch and obtain the time-frequency Fourier coefficients of each dipole component of the 16102 sources. The amplitude of each source was finally obtained as the Euclidean norm of the magnitude of the three corresponding fourier coefficients and averaged over epochs. This led to one source distribution of time-frequency power map per condition (P1, P2 and P3) and subject, which was then used for subsequent analysis.

### 2.6. Event-related (de)synchronization measures

A baseline was defined as the period of time preceding the onset of the first part of the sentence (P1), from 500 ms to 100 ms prestimulus, i.e., during the display of the fixation cross. Event-related “*enhancement*” or “*synchronization*” (ERS) was defined as post-stimulus power increase in a given frequency band compared to the baseline (Pfurtscheller, 2001). Conversely, event-related “*suppression*” or “*desynchronization*” (ERD) was defined as a power decrease (Pfurtscheller, 2001). This baseline was chosen to measure neural processes unfolding as the sentence progressed, including sustained language-related activity, in distinction, e.g., with Hultén et al. (2019) who focused on the effect of the previous context on upcoming stimuli.

In practice, source-level ERS/ERD measures were obtained for each frequency bin by dividing its value with the mean baseline value at the same frequency and subtracting 1.

### 2.7. Statistical analyses

Group-level statistical analysis of the trial-averaged data was performed with a maximum *t* statistic procedure (Blair & Karniski, 1993), which provides a rigorous control of the family-wise error (FWE) rate in the context of multidimensional MEG data (16102 sources x 45 frequency bins x 60 time points), fraught with correlations among neighbouring measurements, while limiting the loss of statistical power (Groppe et al., 2011). For each sentence part (P1, P2 and P3), we started by computing a one-sample *t* statistic for each source-time-frequency point of our dataset. This corresponds to a mass-univariate test against the null hypothesis that conditions induce no power changes compared to the baseline, i.e., no ERS or ERD. The maximum absolute *t* value across this dataset defined the maximum statistic. Its null distribution was generated non-parametrically by recomputing 2000 times a similar statistic after changing the sign of the relative amplitude change data in a random selection of subjects. This amounts to the fact that ERS and ERD are exchangeable under the null. The 95th percentile of this null distribution of maximum was used to establish a significance threshold at a 95% confidence level, fully corrected. All supra-threshold values were deemed significant.

### 2.8. Global spectro-temporal neural dynamics of SC

In order to obtain an overview of the spectro-temporal neural dynamics of the three constituent parts of the SC paradigm, a global time-frequency analysis map was constructed as the mean *t*-value across all sources. Then, only the supra-threshold *t*-values were included to locate the time-frequency points disclosing significant ERS (positive values) or ERD (negative values).

### 2.9. Frequency band and time-resolved mapping associated with SC

The source location associated with supra-threshold power modulations were projected onto a standard MNI brain and visualized with Mricron (Rorden & Brett, 2000; RRID:SCR_002403) to create a frequency and time-resolved functional mapping. This source-space analysis was based on the single statistical threshold obtained from the unique analysis considering all time (60) and frequency (45) points for each of the three sentence parts. Given that a large number of such maps may be built (60 time points x 45 frequency bins), we considered several averages within sliding frequency bands and time windows, allowing for the exploration of ERS/ERD dynamics on different spectro-temporal scales. We first investigated spectrally-resolved maps averaged over the whole post-stimulus period (0–3000 ms) and over classical frequency bands (theta: 4-7 Hz; alpha: 8-11 Hz; low-beta: 12-20 Hz; high-beta: 21-30 Hz; low-gamma: 31-45 Hz). Their temporal development was assessed by averaging over the whole frequency spectrum (1-45 Hz) within five consecutive, post-stimulus, non-overlapping 400 ms-long windows, followed by a last longer window of 1000 ms. This rather coarse temporal segmentation was chosen based on the global time-frequency graph (see Results).

A detailed list of statistically significant local maxima was created for the band-limited maps for further analyses. These local maxima were identified on the thresholded maps further smoothed with an isotropic gaussian kernel (full width at half maximum = 8 mm). Local maxima were discarded if they did not fall within the cortical parcels of the Automated Anatomical Labelling atlas (AAL; Tzourio-Mazoyer et al., 2002; RRID: SCR_003550) and the cerebellar hemispheres to capture plausible cortical activity.

Additionally, to quantify the hemispheric dominance of each frequency band and time window, a laterality index (LI) was extracted from the corresponding maps as the relative difference between the number L of supra-threshold sources in the left hemisphere and the number R of supra-threshold sources in the right hemisphere, i.e., *LI* = (*L* − *R*)/(*L* + *R*). A positive LI thus indicates a leftward dominance, with the maximum value LI = 1 reached if significant sources occur only in the left hemisphere. Likewise, negative LI indicates rightward dominance and the minimum LI = -1 is reached if significant effects arise in the right hemisphere only. A separate LI was calculated for ERS and ERD.

### 2.10 Spectro-temporal neural dynamics in the right vs. left ventral language stream during SC

A region of interest (ROI) covering the ventral language stream was constructed on the basis of a meta-analytic language map (association test map; *p*<.01 corrected for false discovery rate; 1101 studies included; methodological details available on http://neurosynth.org) obtained with the Neurosynth online tool (Yarkoni et al., 2011; RRID:SCR_006798). The map was smoothed with an isotropic Gaussian kernel (full width at half maximum = 10 mm) to account for the difference in location between fMRI activations and MEG sources, which is of the order of the centimetre (e.g., Stippich et al., 2007). A subset of the smoothed and binarized (z-score>0) left-sided map belonging to the posterior middle/inferior temporal (Hickok & Poeppel, 2007) and fusiform (Saur et al., 2008) AAL parcellations (Tzourio-Mazoyer et al., 2002; RRID: SCR_003550) with MNI *y* coordinates between -32 mm (posterior edge of Heschl’s gyrus parcellation) and -76 mm (posterior edge of middle temporal gyrus) was defined as the left part of the ventral stream. Its mirror image constituted the right-sided counterpart. The two ROIs are represented in Figure 5 (rightmost column).

Time-frequency values spatially-averaged within ROI were used to probe neural oscillatory modulations potentially hidden by the stringent statistical threshold imposed by the whole-brain analysis (i.e., loss of sensitivity due to the use of maximum statistics and the large amount of comparisons), and to perform pairwise comparisons of the right vs. left ROIs.

Statistical testing of this difference was based on a maximum statistics procedure similar to the whole-brain analysis but spatially restricted to the ROI. Here, univariate *t*-statistics corresponded to two-sample paired *t*-tests and the null distribution was derived by randomly permuting the labels ‘left’ and ‘right’ among subjects before computing these t-values and extracting their maximum across time, frequencies, and sources in the ROI (number of permutations: 2000).

### 2.11. Data availability statement

The MEG data used in this study will be made available upon reasonable request to the corresponding author and after acceptance by institutional authorities (CUB Hôpital Erasme and Université libre de Bruxelles).

## 3. Results

### 3.1. Global spectro-temporal neural dynamics of SC

The general pattern of the spectro-temporal neural dynamics of SC was visualized using whole-brain-averaged time-frequency maps of ERS/ERD (Figure 2). It consisted of an immediate post-stimulus ERS in the lower frequency range (mainly < 10 Hz) at the onset of each sentence part, though only the initial ERS in P1 in the theta/alpha range (5-8 Hz) reached significance for a short period of time (0-300 ms). This ERS was followed by an ERD peaking around 400 ms mainly in the alpha and beta frequency bands. Significant ERD took place from 250 ms to 600 ms in P1, between 350 ms and the rest of the post-stimulus period in P2 and essentially between 300 ms and 1800 ms in P3. The frequency range of significant ERD was more limited in P1 (alpha and high beta bands) while it stretched from theta to low gamma bands in P2 and P3.

**Figure 2:**
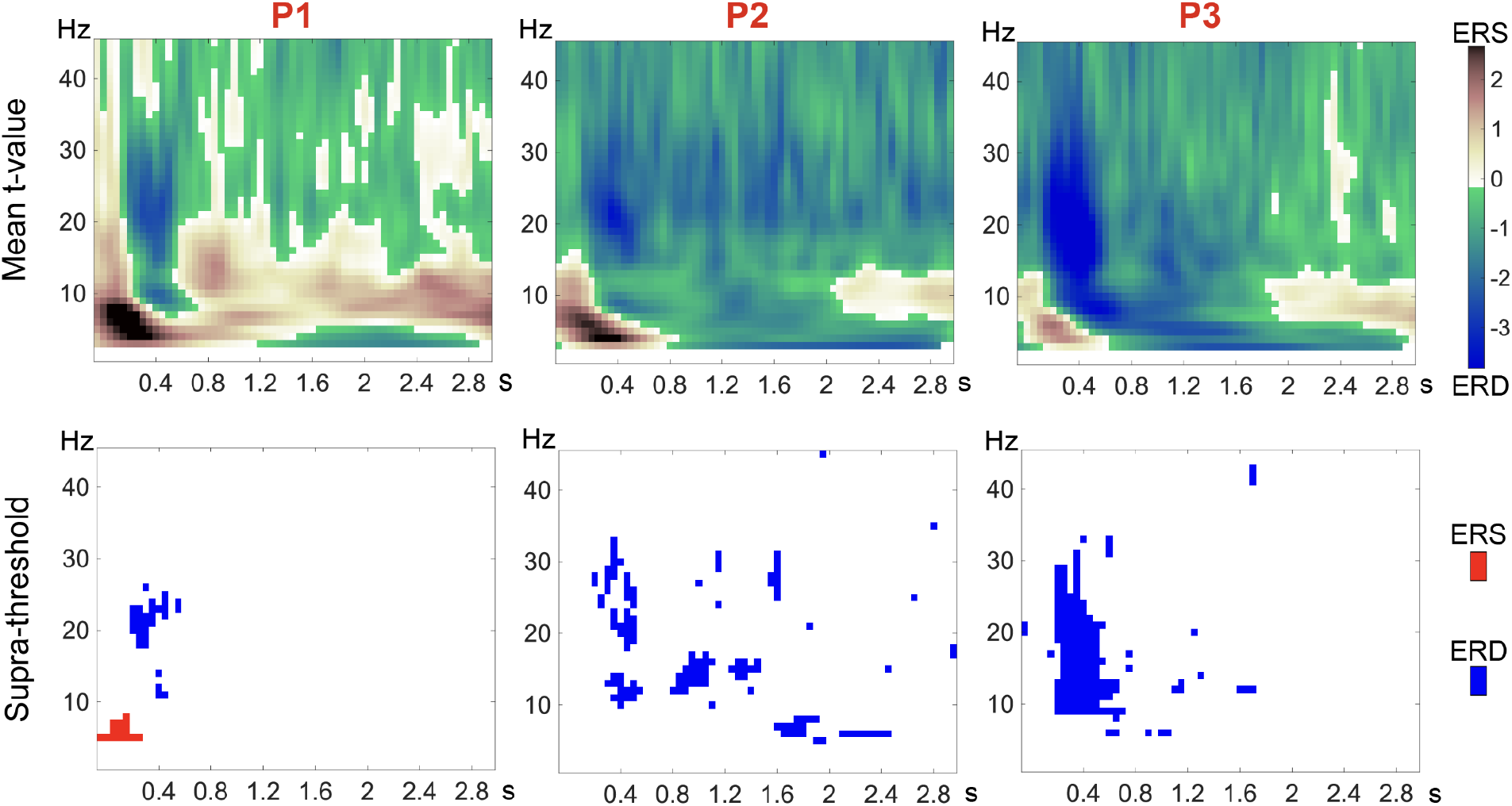
Time-frequency maps for the three parts of the SC paradigm. Whole-brain (average across all sources) power modulations associated with the first (**P1**, noun and determiner; **Left**), second (**P2**, part of predicate; **Middle**) and third part (**P3**, to be completed; **Right**) of the sentence. **Top**. Average of *t*-values of the statmax procedure without statistical masking, with a pink to dark red scale grading for positive (ERS) values and green to dark blue scale grading for negative (ERD) values. **Bottom**. Time-frequency bins of significant (*p*<.05 FWE corrected) ERS (red) or ERD (blue).

Given the temporal spread of the significant ERD in P2 and P3, five non-overlapping 400 ms time windows were selected to cover the first 2 s of the post-stimulus period, followed by a final longer 1000 ms window to study the last second that was only relevant in P2.

### 3.2. Frequency band and time-resolved mapping associated with SC

For each sentence part, brain maps locating significant ERS and ERD in classic frequency bands (theta: 4-7 Hz; alpha: 8-11 Hz; low-beta: 12-20 Hz; high-beta: 21-30 Hz; low-gamma: 31-45 Hz) at any post-stimulus time are represented on Figure 3 along with the corresponding LI measuring the left-right hemispheric asymmetry of significant ERS or ERD. Their temporal development (in five consecutive 400 ms-long windows, followed by a last longer window of 1000 ms) are shown on Figure 4. Associated local maxima are listed in Supplementary Tables 2 and 3.

**Figure 3:**
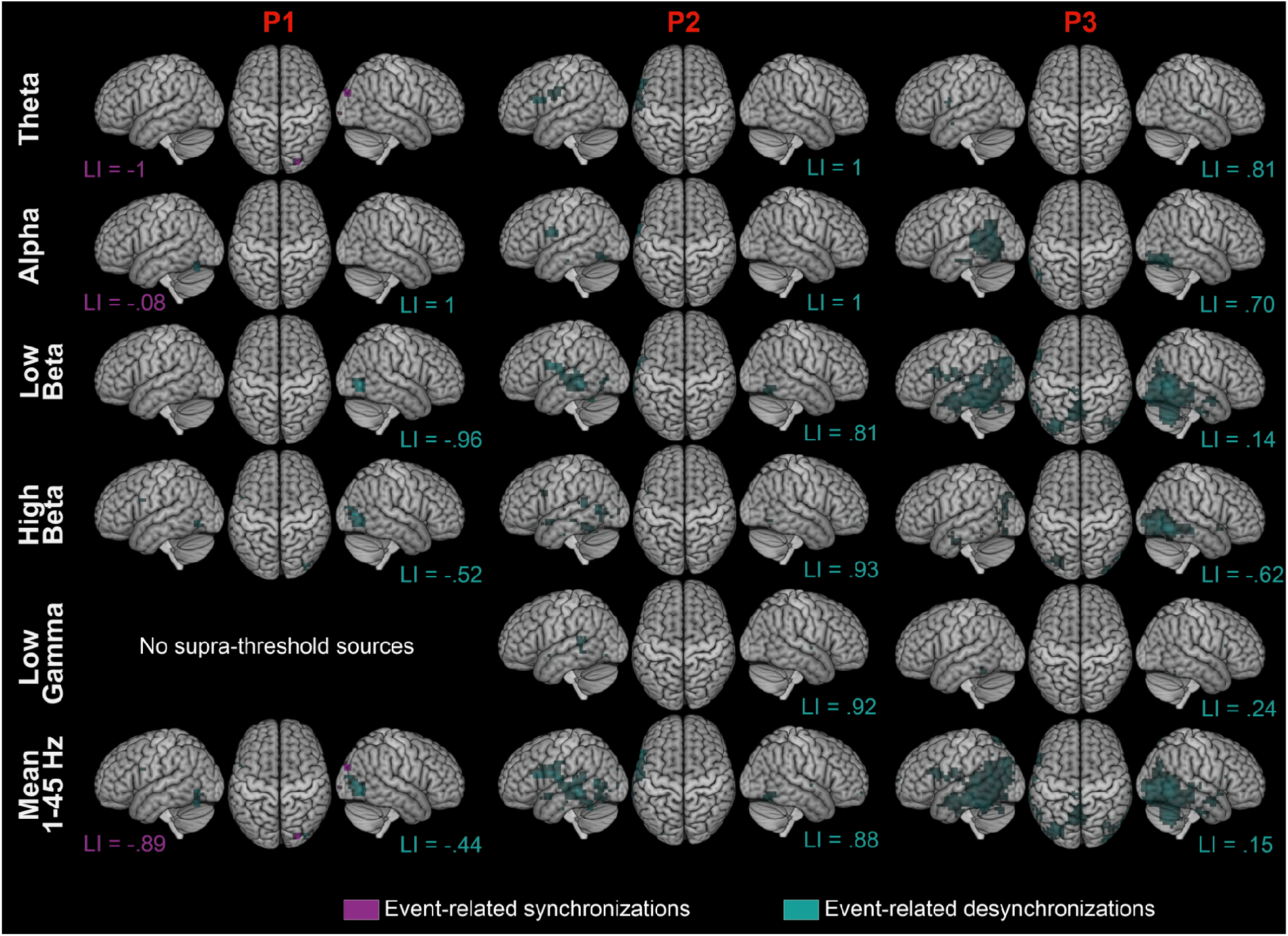
Brain maps showing frequency-dependent changes in neural activity during the three parts of SC. Group-level maps of significant (*p*<.05 FWE corrected) ERS (magenta) and ERD (cyan) averaged across the whole post-stimulus period are superimposed on surface projections (left lateral, top, right lateral) of the standard MNI brain. The supratentorial lateralization indices (LI) derived by counting significant ERS voxels (magenta font) and significant ERD voxels (cyan font) are also indicated (bottom of left and right projections). The first (**P1**, noun and determiner; **Left**), second (**P2**, part of predicate; **Middle**) and third (**P3**, to be completed; **Right**) parts are depicted in the corresponding three columns. Rows correspond to frequency bands in which significant ERS/ERD values were further averaged: theta (4-7 Hz), alpha (8-11 Hz), low beta (12-20 Hz), high beta (21-30 Hz), low gamma (30-45 Hz) and broadband average across the whole frequency range (1-45 Hz).

**Figure 4:**
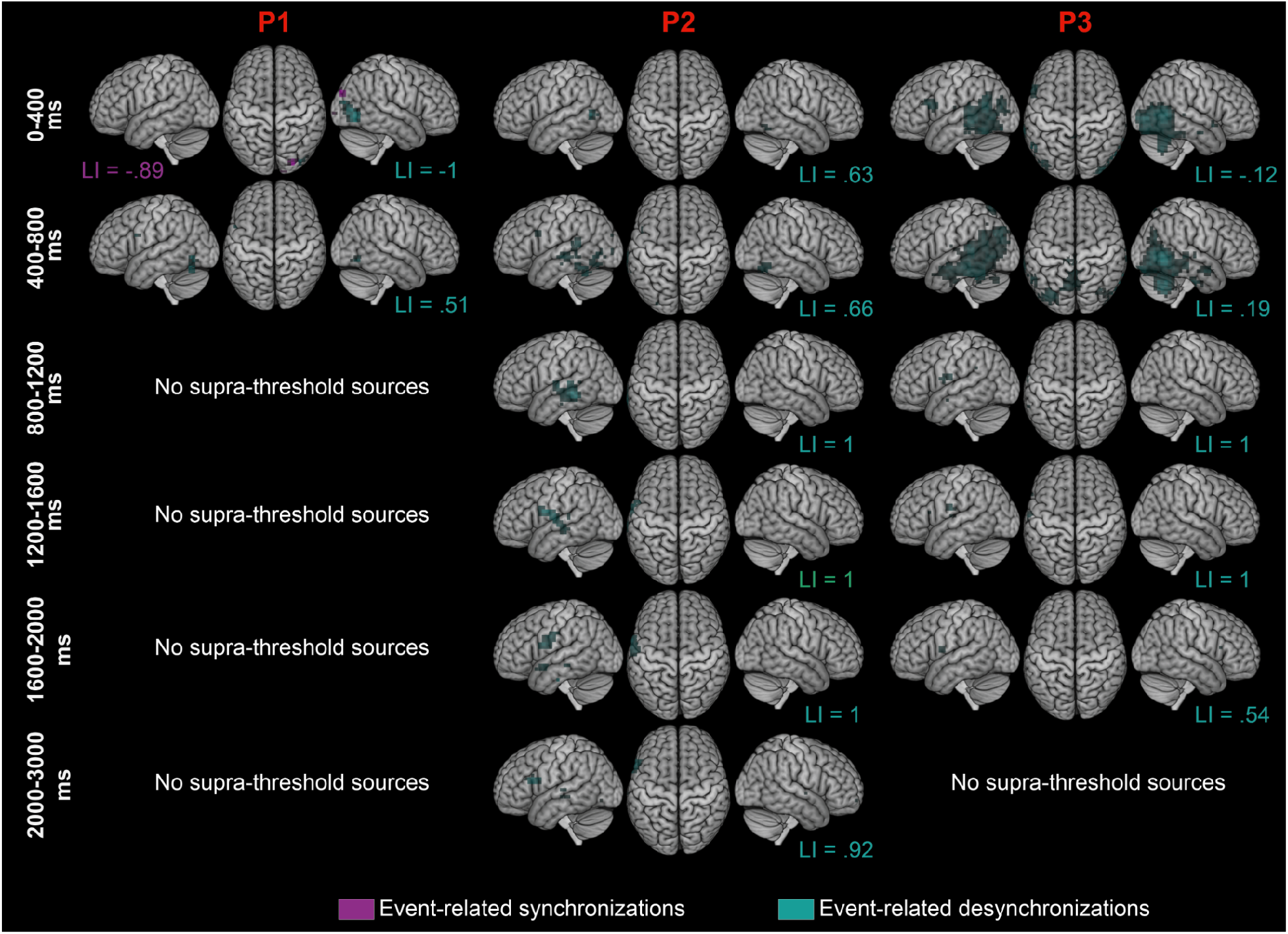
Brain maps showing time-dependent changes in neural activity during the three parts of SC. Group-level significant (*p*<.05 FWE corrected) ERS (magenta) and ERD (cyan) averaged across the whole frequency range (1-45 Hz) superimposed on surface projections (left lateral, top, right lateral) of the standard MNI brain. The supratentorial lateralization indices (LI) derived by counting significant ERS voxels (magenta font) and significant ERD voxels (cyan font) are indicated (bottom of left and right projections). The first (**P1**: noun and determiner; **Left**), second (**P2**, part of predicate; **Middle**) and third (**P3**, to be completed; **Right**) parts are depicted in the corresponding three columns. The rows correspond to the temporal averages of ERS/ERD within consecutive 400 ms-long windows except for the last row (over the last second of the post-stimulus period).

**Figure 5:**
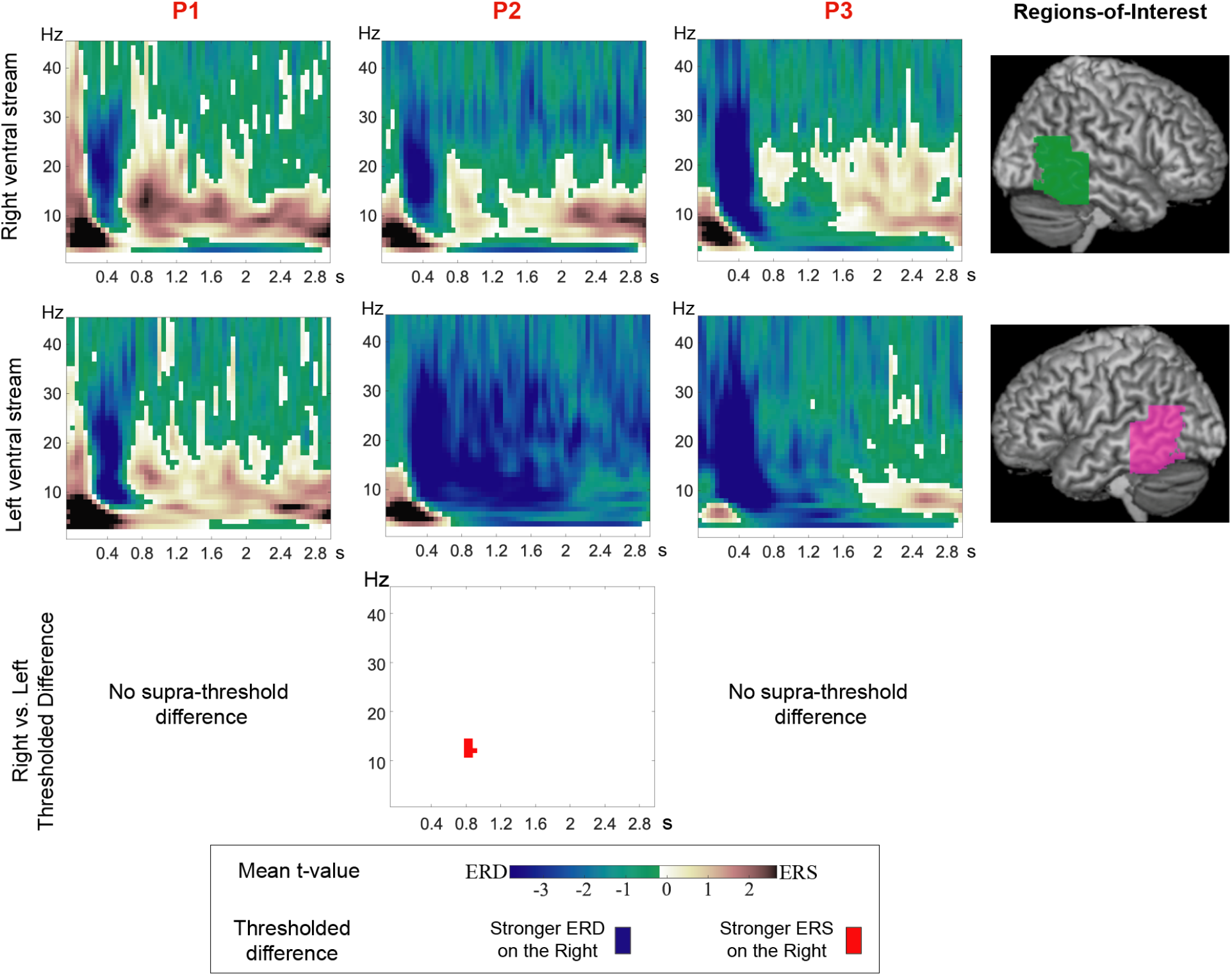
Right vs. left comparative spectro-temporal neural dynamics in the ventral language stream during SC. The first (**P1**: noun and determiner), second (**P2**, part of predicate) and third (**P3**, to be completed) parts are depicted in the corresponding three columns. Time-frequency group-level *t*-statistics (without statistical masking) averaged within a ROI (**Region-of-Interest**; **rightmost column**) are shown in a pink to dark red scale grading the positive (ERS) and green to dark blue scale grading the negative (ERD) *t*-values. **Top**. In the right component of the langage ventral stream (green ROI). **Middle**. In its left component (pink ROI). **Bottom**. Pairwise right vs. left comparison pairs (*p*<.05 with small-volume FWE correction and additional Bonferroni correction for the 3 comparisons considered), coded in red for a stronger ERD or conversely, in blue for a stronger ERD on the right.

During the first part of the SC paradigm (P1; Figs. 3 and 4, left column), the first theta and alpha ERS occurring in the 0-400 ms time window were located in bilateral calcarine cortices (not visible on source projections but detailed in Supplementary Table 1-2) as well as in the right lateral occipital cortex, leading to a right-dominant LI. The low-beta part of the ERD occurring in the same time window appeared in right lateral occipito-temporal and posterior fusiform regions, also leading to a right-dominant LI. Postero-inferior ERD became more bilateral and supported by high beta in the 400-800 ms time-window, which also disclosed a small cluster of high-beta ERD in the left inferior frontal gyrus (IFG), leading to a left-dominant LI. No ERS/ERD was significant after 800 ms.

During the second part (P2; Figs. 3 and 4, middle column), the 0-800 ms time windows displayed bilateral beta ERD in the fusiform gyri with leftward dominance, along with left-sided beta ERD in the lateral occipito-temporal cortex and in the IFG. In the 800-3000 ms time windows, broadband ERD were located in the lateral left temporal cortex (alpha, beta and low-gamma) and in left IFG and rolandic opercular regions (theta, alpha and beta; 1200-3000 ms time-windows), as well as the left posterior cingulate cortex (PCC; high-beta and low-gamma; 1200-1600 ms time-window; not visible on the projections). These ERD led to left-dominant LI for each time-window and frequency range.

The third part (P3; Figs. 3 and 4, right column) presented stronger and more extended ERD located in bilateral temporo-occipito-parietal areas, including more anterior aspects of the lateral temporal cortices. They occurred mostly in the 0-800 ms time windows and involved theta, alpha, beta and gamma frequency bands. In the 400-800 ms time window, beta-band ERD also emerged in the left posterior cingulate cortex (PCC) and precuneus, middle occipital gyri and angular gyri bilaterally and in the superior part of the cerebellum. Moreover, theta and beta ERD were observed in the left IFG and rolandic operculum from 0 to approximately 2000 ms post-stimulus. Finally, a late (1600-2000 ms) and small gamma-band ERD was also present in the right IFG (not visible on the figure as it involved only a few brain sources). These ERDs led to left-dominant LI, except for the 0-800 ms time windows and beta and gamma-bands.

Overall, in each sentence part, the alpha and beta frequency bands (Fig. 3, rows 2-4) accounted for the majority of the significant ERD in the wide-band, temporally averaged maps (Fig. 3, last row).

### 3.3. Right vs. left comparative spectro-temporal neural dynamics in the ventral language stream during SC

Among the three pairwise comparisons between the right and left components of the ventral language stream, the only significant (*p*=.0495, with small-volume FWE and Bonferroni correction) difference emerged in P2 in the alpha/low-beta bands (11-14 Hz) in a cluster centered on 900-1050 ms. This difference originated from clearly more ERD on the left, with a sustained 8-30 Hz ERD after the initial dominant ERD centered on 400 ms, contrasting with a more subtle 8-20 Hz ERS trail on the right.

Infra-threshold analysis did not demonstrate prolonged ERD on the right side compared to the left in P1 or P3. Indeed, in P1, the time-frequency dynamics was similar to the global spectro-temporal dynamics of SC on both sides. In P3, the ERD centered on 400 ms was actually slightly narrower.

## 4. Discussion

This MEG study demonstrates that SC elicits three successive significant neural response patterns characterized by distinct spectro-temporal signatures and anatomical locations for each sentence part: (i) an early (<300 ms post-stimulus) bilateral 4-10 Hz ERS in occipital cortices that is most conspicuous in P1; (ii) an 8-30 Hz ERD at about 400 ms post-stimulus involving the ventral language stream of both hemispheres as well as posterior nodes of the default mode network (DMN); and (iii) a late (> 800 ms post-stimulus) 8-30 Hz ERD involving the left dorsal language stream observed in P2 and P3. Furthermore, the left ventral language stream displayed prolonged 8-30 Hz ERD from 800 ms post-stimulus during P2 compared to its right homolog which showed an 8-20 Hz ERS from 800 ms post-stimulus.

### 4.1 First neural response pattern of SC: low-level visual processing

The first neural response associated with SC was characterized by a 4-10 Hz ERS which was localized in the bilateral primary visual cortices and the right superior occipital gyrus during P1. A similar response was also observed in P2 and P3, but it did not reach statistical significance. This response is attributed to the visual processing of written stimuli given its rapid post-stimulus (0-300 ms) occurrence. This first response is in agreement with the timing and topography of previously reported MEG responses at play during reading and that are related to non-specific, pre-lexico-semantic visual features (Pammer et al., 2004; Tarkiainen et al., 1999; for a review, see Salmelin 2007).

### 4.2. Second neural response pattern of SC: bihemispheric semantic processing

The second response was associated with an ERD in the alpha-beta (8-30 Hz) frequency range centered at about 400 ms post-stimulus (starting at about 200-300 ms post-stimulus and lasting 300 to 400 ms) in each part of the sentence, but most prominent during P2 and P3. It was located in the bilateral posterior inferior/middle temporal gyri, and also in the left IFG during P2.

The timing of this response probably relates to the previously described N400m evoked response associated with lexico-semantic processing (Halgren et al., 2002; Salmelin, 2007). As in the present study, this response has also been shown to involve frontal brain areas, co-occurring as early as from 200-250 ms (Halgren et al., 2002; Helenius, 1998; Pammer et al., 2004; Vartiainen et al., 2011). In accordance with previous studies (Halgren et al., 2002; Hultén et al., 2019; Pammer et al., 2004), the observed bilateral occipito-temporal ERD and the co-modulations occurring around 400 ms in the left inferior frontal (in P1,2,3) and the temporal (left in P2 and bilateral in P3) areas are jointly attributed to the same N400m response of SC, where phono-lexico-semantic integration concomitantly occurs with the processing of visual words to conduct the reading process effectively.

The bi-hemispheric N400m ERD in the posterior temporal lobes consistently colocalized with the ventral stream of language (Hickok & Poeppel, 2007). It has been suggested that word- and sentence-level comprehension rely on different processing strategies in both hemispheres (Federmeier et al., 2008) and that specific right temporal contribution is involved in sentence-level processing (Vigneau et al., 2010). However, at this stage we did not find significant interhemispheric asymmetries in the spectral signature that could suggest a differential contribution.

In addition to the involvement of the ventral language stream, posterior parasagittal and lateral parietal cortices also presented ERD in the beta band at about 400 ms post-stimulus, which is consistent with the concept of a distributed semantic network (Binder, 2015; Graves, 2010; Paunov et al., 2019; Seghier & Price, 2012; Whitney et al., 2009), involved in the semantic processes of sentences (as reviewed by Weiss & Mueller, 2012). Contrastingly, in fMRI, Graves (2010) and Whitney et al. (2009) relied on semantic tasks being contrasted with other high-level language tasks (e.g., familiar highly meaningful phrases vs. unfamiliar phrases with minimal meaning) to demonstrate a relative increase in fMRI signal in the regions classically attributed to the default mode network (for a review, see e.g., Raichle, 2015). These nodes are typically anticorrelated with the task at hand and Seghier & Price (2012) found that their degree of deactivation was dependent on the semantic nature of the task. We here find a common beta-band ERD signature throughout the whole extended semantic network, in distinction with the known heterogeneity of its fMRI counterpart (activations and deactivations). The engagement of the posterior DMN may facilitate the retrieval and integration of relevant informational and contextual content (Buckner et al., 2008) in order to select a relevant completion.

Of note, the beta-band ERD in the anterior lobe of the right cerebellum in P3 could support the potential semantic role of the cerebellum in semantic language processing. Right cerebellar contribution to the language function has been largely documented (for reviews, see, e.g., Mariën et al., 2013; Price, 2012) and even used as a crossed cerebro-cerebellar lateralization aid in a presurgical functional language mapping perspective (Méndez Orellana et al., 2015). However, given the many language subprocesses it has been associated with (Mariën et al., 2013), the ERD in the right cerebellum might reflect its role as a general contributor/modulator of higher cognitive processes rather than representing another specific node of the distributed semantic network.

### 4.3. Third neural response pattern of SC: left-sided dorsal stream integration

The third and last neural response pattern was characterized by an alpha-beta ERD in P2 and P3, taking place from >800 ms post-stimulus. Contrary to previous responses, this ERD, most clearly demonstrated in P2, presented a clear leftward dominance (LI ERD = .54 to 1) and mainly involved the left fronto-temporal regions. These areas (IFG, superior temporal gyrus) are topographically consistent with the key regions of the classical Broca–Wernicke–Lichtheim–Geschwind model (Tremblay & Dick, 2016) and of the more recently proposed MUC (Memory, Unification, Control) model (Hagoort, 2013). Information processed in the previous N400 step might be integrated during these long-lasting ERD within the left hemisphere, dominant for language processing (Tzourio-Mazoyer et al., 2017), to be further semantically and syntactically unified, and to allow control (Hagoort, 2013) and selection of the required words to properly complete the sentence. They are also anatomically in line with the left-dominant dorsal stream of the dual stream model (Hickok & Poeppel, 2007), including the revisited, speech production Wernicke’s area (Binder, 2015). They might therefore also reflect some level of production and phonological processes (Vigneau et al., 2006), such as verbal working memory to maintain the sentence content (Meltzer et al., 2017).

In the ventral stream, no whole-brain suprathreshold modulation was demonstrated at this stage. However, focused ROI analyses showed a sustained alpha-beta ERD on the left, contrasting with moderate ERS in those frequency bands on the right in P2 and P3 (the right vs. left difference reaching significance only in P2). Considering that alpha-beta bands ERS may be considered as an electrophysiological correlate of a deactivated/inhibited cortical area (for reviews, see, e.g., Klimesch et al., 2007; Pfurtscheller, 2001), we here suggest a left-to-right interhemispheric inhibition (Tzourio-Mazoyer et al., 2017) where the left hemisphere would recover its dominance for efficient left intra-hemispheric integration (Gotts et al., 2013), through a left-to-right “top-down” inhibition by means of alpha-beta modulations (Fries, 2015). Indeed, the genetic, developmental, structural and functional factors (for reviews, see e.g., Stephan et al., 2007; Tzourio-Mazoyer et al., 2017) leading to left hemisphere specialization for language, might position this hemisphere as the actual higher order language hemisphere for subsequent SC process, endowed with the control component of the MUC model (Hagoort, 2013). We therefore did not find signs of sustained engagement of the right hemisphere that could indicate retention of multiple meanings for prolonged periods of time (Kircher et al., 2001) or integration of information over longer timescales as in auditory speech processing (Hickok & Poeppel, 2007).

### 4.4. Neural processing in the ventral and dorsal streams of the language network during SC is dominated by transient alpha-beta frequency band power decrease

Spectral specialization in language processing has been suggested for different frequency bands due to selectivity to certain linguistic manipulations (for a review, see, e.g., Meyer, 2017). It can also be suspected given the observation of a spatio-spectral pattern in the language network during task (Goto et al., 2011) or at rest (Coolen et al., 2020), whereby lower frequency bands (theta, alpha) tend to localize ventrally and posteriorly while higher frequency bands (beta, gamma) are more dorsal and anterior, in their modulations or connectivity.

However, in this study, alpha and beta ERD represented the main neural activity modulations observed within the whole language network. The main alpha-beta-band spectrum in language-related neural processes found here is in agreement with the prominence of the ERD in those two frequency bands reported in previous MEG language studies (Goto et al., 2011; Lam et al., 2016; Piai et al., 2015), and with the concept of a broadband alpha-beta (8-30 Hz) ERD in sentence processing (Kielar et al., 2015; Meltzer et al., 2017; Meltzer & Braun, 2011). The alpha (e.g., Obleser & Weisz, 2012; Wang et al., 2018) and beta (e.g., Piai et al., 2015; Weiss & Mueller, 2012) frequency bands have been shown to be involved in language processes in various experimental designs and to serve as the main channels for interactions within the language network (Schoffelen et al., 2017). Beyond information transfer (Buzsáki & Draguhn, 2004), the observed coordinated alpha-beta ERD constituting the response patterns in the extended semantic network and in the dorsal stream of language might also reflect the formation of specific dynamic workspaces (Lopes da Silva, 2013), for efficient and sustained language processing. In addition, a spatial correlation was previously shown between task-related ERD in alpha-beta bands and the fMRI task-related changes (Hall et al., 2014; Mukamel, 2005), particularly in language tasks (Singh et al., 2002).

### 4.5. Limitations

In contrast with some previous sentence-level paradigms (e.g., Kielar et al., 2015; Meltzer et al., 2017), this study did not use specific linguistic manipulations. In particular, the lack of controlled semantic modulations (e.g., theory-of-mind, semantic complexity) did not allow for the refinement of the relative contribution of the constituent parts of the observed extended semantic network, especially that of the posterior nodes of the DMN. We also did not use standardized sentences (e.g., Kircher et al., 2001; Wilson et al., 2017) nor recorded the completions given by the subjects. Moreover, our design (3 second-long variable length stimuli) did not allow for a fine-grained temporal analysis (e.g., Halgren et al., 2002; Hultén et al., 2019). We instead provided a continuous, detailed neuromagnetic overview of the neural dynamics occurring during a classic nonstandardized SC paradigm.

Finally, the rather limited population, the prior-free whole-brain analysis and the stringent maximum statistics that we used might have impeded statistical power. For instance, the immediate post-stimulus lower-level visual processing step (described in 4.1) reached statistical significance only in P1. In the other two parts of the sentence (P2, P3), this approach might be solely picking up the maximum difference from baseline driven by the language-related ERD. This probably includes the temporal modulations around 400 ms associated with the increasing semantic load and N400m effect (Halgren et al., 2002), that remained infra-threshold in P1. Similarly, the relative lack of significant deviations from baseline observed in the theta and gamma bands could be partly due to the biased sensitivity of our statistical approach to the stronger and most consistent ERD in alpha and beta bands. Conversely, the spectro-temporal modulations we reported are highly likely to represent true neural responses. The use of stringent maximum statistics also ensures the robustness of the observed neural activity modulations.

## 5. Conclusions

This study elucidates the neural dynamics that accompany the SC process, consisting of three successive neural response patterns associated with distinct spatial and spectro-temporal characteristics: an early low-level visual response; an intermediate bihemispheric semantic processing within an extended semantic network; a late left-hemisphere dorsal language stream integration.

We brought critical insight into the differential contribution of the ventral language stream, where the right-sided contribution is more transient and associated with some signs of subsequent inhibition. Furthermore, we demonstrated a common fingerprint throughout an extended semantic network. This shared signature is in contradistinction with its known heterogeneous hemodynamics (i.e., task-correlation of fMRI signal in temporal areas versus anticorrelation in the DMN). Therefore, this study may encourage more MEG research in sentence-level paradigms and their engagement of the extended semantic network. MEG-fMRI correspondence studies are also needed to elucidate the apparent heterogeneity of the neurovascular coupling within that network.

This adaptation of the fMRI SC paradigm to MEG paves the way for neuromagnetic preoperative language mapping in neurosurgical patients, free of neurovascular coupling issues (see e.g., Pak et al., 2017). Thanks to the dynamic dimension brought by MEG in SC, this may help in the identification of regions that need to be preserved among all activated regions disclosed by the static mapping fMRI provides. It also opens new perspectives in understanding the physiopathology of language alterations in various brain disorders. For instance, sentence completion tests have been administered to evaluate cognition in neurodegenerative diseases, such as Alzheimer’s disease (Kim & Thompson, 2004; Martyr et al., 2019) Parkinson’s disease (Martyr et al., 2019; Siquier & Andrés, 2021) and amyotrophic lateral sclerosis (Abrahams et al., 2005). MEG should bring novel insights in the spectro-temporal correlates of previously reported abnormalities.

## CRediT author statement

**T. Coolen**: Conceptualization, Software, Formal Analysis, Investigation, Writing - Original Draft, Visualization. **A**.**M. Dumitrescu**: Conceptualization, Formal Analysis, Investigation, Writing - Review and Editing, Visualization. **V. Wens**: Methodology, Software, Formal Analysis, Resources, Writing - Review and Editing. **M. Bourguignon**: Software, Review and Editing. **G**.**L. Gómez**: Methodology, Software, Resources. **A. Rovai**: Software, Resources. **N. Sadeghi**: Writing - Review and Editing, Supervision. **C. Urbain**: Writing - Review and Editing, Supervision. **S. Goldman**: Writing - Review and Editing, Supervision, Project Administration, Funding Acquisition. **X. De Tiège**: Conceptualization, Writing - Review and Editing, Project Administration, Funding Acquisition.

## Acknowledgements

Tim Coolen (MD, PhD student) is Clinical Master Specialist Applicant to a PhD at the Fonds de la Recherche Scientifique (FRS-FNRS, Brussels, Belgium). Alexandru Dumitrescu is supported by the Fonds Erasme (Research Convention “Les Voies du Savoir”, Brussels, Belgium). Gustavo Lucena Gómez was supported by the Excellence of Science “MEMODYN” of the Fonds de la Recherche Scientifique (FRS-FNRS, Brussels, Belgium). Xavier De Tiège is a Postdoctorate Clinical Master Specialist at the FRS-FNRS (Brussels, Belgium). This study and the MEG project at the CUB Hôpital Erasme are financially supported by the Fonds Erasme (Research Convention “Les Voies du Savoir”, Brussels, Belgium). Funding sources had no involvement in the present study. Declarations of interest: none.

We extend our gratitude to Thomas Amighi (Master Student at the Faculty of Psychology, Université libre de Bruxelles) for collecting the behavioral data presented in the Supplementary Materials.

