## Supplementary for "Spectro-temporal neural dynamics during sentence completion"

### Supplementary Material

#### Supplementary Table 1. Sentences to complete

| | Part 1: subject | Part 2: part of predicate | English translation |
|---|---|---|---|
| 1 | La poule | pond un | The hen lays an |
| 2 | La grand-mère | raconte une | The grandmother tells a |
| 3 | Cet homme | rase sa | This man shaves his |
| 4 | Les chats | attrapent les | Cats catch |
| 5 | La vache | donne du | The cow gives us |
| 6 | La capitale | française est | The French capital is |
| 7 | Les chiens | rongent un | The dogs gnaw a |
| 8 | Le verre | tombe par | The glass falls to the |
| 9 | Une image | vaut mieux que mille | A picture is worth a thousand |
| 10 | La secrétaire | écrit une | The secretary writes a |
| 11 | Le singe | mange une | The monkey eats a |
| 12 | La petite fille | joue avec une | The little girl plays with a |
| 13 | Les déchets | se jettent dans | Litter is thrown in the |
| 14 | Les girafes | ont un long | Giraffes have a long |
| 15 | Les lapins | ont des grandes | Rabbits have big |
| 16 | Les dentistes | soignent les | The dentists treat |
| 17 | Le rire | est le propre de | To laugh is proper to |
| 18 | Le bonnet | se porte sur | The hat is worn on the |
| 19 | Les lunettes | améliorent la | Glasses improve |
| 20 | Le boucher | vend de la | A butcher sells |
| 21 | Les vampires | sucent du | Vampires suck |
| 22 | Les avions | volent dans le | Planes fly in the |
| 23 | Le métro | est un moyen de | The metro is a means of |
| 24 | Cette armoire | est fermée à | This cabinet is locked with a |
| 25 | Les absents | ont toujours | Absent is always in the |

| | | | |
|---|---|---|---|
| 26 | L’union | fait la | Unity is |
| 27 | Le fleuriste | a arrosé les | The florist watered the |
| 28 | Cet étudiant | est le meilleur de sa | This student is the best of his |
| 29 | La voiture | est tombée en | The car broke |
| 30 | Un tien | vaut mieux que | A bird in the hand is worth two in |
| 31 | Toutes les bonnes choses | ont une | All good things come to an |
| 32 | La cigarette | s'écrase dans le | Cigarettes are put out in the |
| 33 | Les trains | roulent sur des | Trains run on |
| 34 | Le boulanger | vend du | A baker sells |
| 35 | L’horloge | indique l’ | A clock shows the |
| 36 | Demain | est un autre | Tomorrow is another |
| 37 | Le chat | est vite monté sur | The cat quickly jumped on |
| 38 | La police | a arrêté | The police arrested |
| 39 | Ce gamin | a beaucoup joué | This kid played a lot |
| 40 | Le cuisinier | a préparé | The cook prepared |
| 41 | Cet employé | va nettoyer | This employee will clean |
| 42 | Les oiseaux | se sont posés | The birds have landed |
| 43 | Les vacanciers | aiment visiter | Vacationers like to visit |
| 44 | Ces hommes | travaillent | These men work |
| 45 | Les invités | sont tous venus | The guests all came |
| 46 | La femme | a souvent porté | The women often wore |
| 47 | Cette entrée | était bloquée | This entrance was blocked |
| 48 | Le printemps | est la saison | Spring is the season of |
| 49 | Notre fils | bâillait | Our son was yawning |
| 50 | Le chien | aboyait toujours | The dog always barked |
| 51 | Sa marraine | lui a acheté | His godmother bought him |
| 52 | Cette pièce | contient un grand | This room contains a big |
| 53 | La fleur rouge | se trouve | The red flower is |
| 54 | Le client | règle ses achats | The customer pays his purchases |
| 55 | La fenêtre | donne sur | This window opens on |

| | | | |
|---|---|---|---|
| 56 | Cet homme | est arrivé enfin | This man has finally arrived |
| 57 | Sa fille | est plus grande | His daughter is taller |
| 58 | La directrice | a convoqué | The director convened |
| 59 | Ses affaires | sont rangées | His belongings are stored |
| 60 | La cave | était remplie | The basement was filled |
| 61 | Les lions | dévoraient | Lions were devouring |
| 62 | Ce magasin | se spécialise dans | This shop specializes in |
| 63 | La guerre | a détruit les | The war destroyed |
| 64 | La discussion | était portée sur | The discussion was about |
| 65 | La réunion | était seulement | The meeting was only |
| 66 | Les intempéries | ont causé | Bad weather caused |
| 67 | Ce musicien | joue très bien | This musician excels at |
| 68 | La caisse | était remplie de | The box was filled of |
| 69 | Le détective | a trouvé | The detective found |
| 70 | Les enfants | se sont promenés | The children went for a walk |
| 71 | Cet élève | a dessiné | This pupil drew |
| 72 | Le ballon | est tombé | The ball fell |

The cloze probability was assessed for each sentence as in Bloom & Fischler (1980), except that only one noun conveying the main idea was taken into consideration if the answer contained more than one word. Half of the sentences (1-36) were composed to have a high cloze probability (HC) and the other half (37-72) a low cloze probability (LC). The two samples were assessed behaviorally on an independent population of 23 undergraduate students of the Faculty of Psychology of the Université libre de Bruxelles (Brussels, Belgium) to ensure the HC sentences (mean cloze probability and standard deviation: 83 +/- 18 %) were significantly different (two-sample t-test; $p<.001$) from the LC sentences (24 +/- 10 %). An equal number of LC and HC was randomly picked for each subject.

*Bloom, P. A., & Fischler, I. (1980). Completion norms for 329 sentence contexts. Memory & Cognition, 8(6), 631–642. https://doi.org/10.3758/BF03213783*

**Supplementary Table 2: local maxima of frequency-dependent changes**

| | P1 | | | | | P2 | | | | | P3 | | | | |
|---|---|---|---|---|---|---|---|---|---|---|---|---|---|---|---|
| **Band** | **MNI** | | | **Location** | **t-value** | **MNI** | | | **Location** | **t-value** | **MNI** | | | **Location** | **t-value** |
| **Theta** | 25 | -89 | 28 | SOG(R) | 11.54 | -57 | 19 | 18 | IFGtr | -11.12 | 71 | -17 | 2 | STGa(R) | -10.52 |
| | 14 | -92 | 3 | CAL(R) | 10.07 | -60 | -7 | 28 | ROLop | -10.31 | -52 | 1 | 15 | ROLop | -10.35 |
| | | | | | | -35 | -5 | 15 | ROLop | -9.91 | -55 | -7 | -16 | MTGa | -9.65 |
| **Alpha** | 1 | -67 | 10 | CAL | 10.11 | -61 | 2 | 18 | ROLop | -10.24 | 49 | -67 | -17 | FUSp(R) | -11.32 |
| | -45 | -69 | -21 | FUSp | -10.66 | -55 | -67 | -16 | ITGp | -10.17 | -52 | -59 | 15 | AG | -11.18 |
| | | | | | | | | | | | -55 | -40 | 5 | MTGm | -11.02 |
| | | | | | | | | | | | -42 | -56 | 4 | MTGp | -10.84 |
| | | | | | | | | | | | -45 | -59 | -3 | MTGp | -10.59 |
| | | | | | | | | | | | -63 | -30 | -22 | ITGm | -9.78 |
| **Low-beta** | 37 | -71 | -4 | FUSp(R) | -11.25 | -58 | -36 | -6 | MTG | -12.23 | 51 | -60 | 1 | MTGp(R) | -13.17 |
| | | | | | | -54 | 3 | 19 | ROLop | -12.11 | -30 | -72 | 29 | MOG/AG | -12.48 |
| | | | | | | -60 | -8 | 16 | ROLop | -10.96 | 38 | -60 | -40 | CBL1(R) | -11.95 |
| | | | | | | -30 | -54 | -26 | FUSm | -10.85 | -57 | -40 | 6 | MTGp | -11.54 |
| | | | | | | 48 | -65 | -17 | ITGp(R) | -10.34 | -58 | -55 | 25 | SMG | -11.43 |
| | | | | | | -32 | -75 | 2 | FUSp | -10.13 | -23 | -16 | -27 | PH | -11.26 |
| | | | | | | -16 | -54 | -19 | FUSm | -10.11 | 56 | -1 | -20 | MTGa(R) | -11.22 |
| | | | | | | -41 | -71 | -4 | ITGp | -9.79 | -49 | -50 | -23 | ITGp | -11.21 |
| | | | | | | -52 | -67 | -16 | ITGp | -9.51 | -56 | -34 | -17 | MTGp | -11.19 |
| | | | | | | | | | | | -61 | -11 | 20 | ROLop | -10.88 |
| | | | | | | | | | | | -27 | -50 | -20 | FUSm | -10.79 |
| | | | | | | | | | | | -44 | 2 | 12 | ROLop | -10.68 |
| | | | | | | | | | | | -55 | -11 | -21 | MTGa | -10.6 |
| | | | | | | | | | | | -3 | -42 | 40 | PCC | -10.57 |
| | | | | | | | | | | | -55 | 16 | 19 | IFGop | -10.5 |
| | | | | | | | | | | | -8 | -66 | 62 | PC | -10.3 |
| | | | | | | | | | | | 68 | -29 | 1 | STGm(R) | -10.19 |
| | | | | | | | | | | | -34 | -73 | -32 | CBL1 | -10.17 |
| | | | | | | | | | | | 41 | -15 | 7 | ROLop(R) | -10.04 |

| | | | | | | | | | | | | | | | |
|---|---|---|---|---|---|---|---|---|---|---|---|---|---|---|---|
| **High-beta** | -29 | -64 | -9 | FUSp | -10.82 | -42 | -61 | 5 | MTGp | -11.96 | 53 | -60 | -9 | ITGp(R) | -13.03 |
| | 37 | -73 | -4 | FUSp(R) | -10.6 | -58 | -45 | 14 | STGp | -10.85 | 28 | -81 | -4 | FUSp(R) | -12.31 |
| | -52 | 8 | 25 | IFGop | -10.15 | -30 | -76 | 2 | FUSp | -10.82 | -31 | -69 | 27 | MOG/AG | -10.64 |
| | -40 | -68 | -9 | FUSp | -9.81 | -43 | 2 | -12 | STGa | -10.38 | 48 | 10 | -10 | STGa(R) | 10.64 |
| | | | | | | -13 | -40 | -17 | FUSm | -10.37 | -33 | -82 | 36 | MOG/AG | -10.57 |
| | | | | | | -62 | -45 | -20 | ITGp | -10.25 | 32 | -43 | -8 | FUSm(R) | -10.41 |
| | | | | | | -67 | -29 | -11 | MTGm | -9.99 | -52 | -12 | -28 | ITGa | -10.21 |
| | | | | | | -22 | -91 | 4 | FUSp | -9.98 | -49 | -79 | 7 | MTGp | -10.2 |
| | | | | | | -45 | -16 | -16 | MTGm | -9.98 | -41 | -80 | 20 | MTGp | -10.09 |
| | | | | | | -7 | -47 | 15 | PCC | -9.98 | 12 | -56 | 45 | PC(R) | -10.07 |
| | | | | | | -48 | 11 | 28 | IFGop | -9.7 | -47 | -78 | -12 | FUSp | -10.05 |
| | | | | | | 49 | -63 | -11 | ITGp(R) | -9.55 | -6 | 73 | 12 | FP | -9.91 |
| | | | | | | -54 | -69 | 1 | MTGp | -9.51 | -43 | -72 | -3 | MTGp | -9.84 |
| **Low-gamma** | *No supra-threshold sources* | | | | | -7 | -47 | 15 | PCC | -10.04 | -64 | -49 | -22 | ITGp | -11.28 |
| | | | | | | -50 | -40 | 8 | MTGp | -9.8 | 51 | 19 | 15 | IFGop(R) | -10.36 |
| | | | | | | 61 | -7 | -1 | STG(R) | -9.73 | 55 | -53 | -26 | ITGp(R) | -9.91 |
| | | | | | | -21 | -86 | 4 | FUSp | -9.66 | -41 | -19 | -24 | ITGm | -9.86 |
| | | | | | | -24 | -83 | 1 | FUSp | -9.63 | | | | | |
| | | | | | | -40 | -77 | -11 | FUSp | -9.57 | | | | | |
| | | | | | | -45 | 4 | -11 | STGa | -9.55 | | | | | |

**Supplementary Table 3: local maxima of time-dependent changes**

| | P1 | | | | | P2 | | | | | P3 | | | | |
|---|---|---|---|---|---|---|---|---|---|---|---|---|---|---|---|
| Time | MNI | | | Location | t-value | MNI | | | Location | t-value | MNI | | | Location | t-value |
| 0-400 ms | 37 | -72 | -4 | FUSp(R) | -11.09 | -42 | -61 | 5 | MTGp | -11.96 | 48 | -60 | 4 | MTGp(R) | -12.56 |
| | 14 | -92 | 3 | CAL(R) | 10.07 | 50 | -62 | -13 | FUSp(R) | -9.55 | 28 | -80 | -4 | FUSp(R) | -12.43 |
| | 1 | -67 | 10 | CAL | 10.11 | | | | | | -52 | -59 | 14 | MTGp | -11.72 |
| | 25 | -89 | 28 | SOG(R) | 11.54 | | | | | | -49 | -57 | -14 | ITGp | -11.39 |
| | | | | | | | | | | | -55 | -41 | 3 | MTGm | -10.76 |
| | | | | | | | | | | | 48 | 10 | -10 | STGa(R) | -10.61 |
| | | | | | | | | | | | -55 | 16 | 19 | IFGop | -10.5 |
| | | | | | | | | | | | -3 | -45 | 37 | PCC | -10.48 |
| | | | | | | | | | | | 41 | -58 | -28 | CBL1(R) | -10.44 |
| | | | | | | | | | | | -38 | -56 | 6 | MTGp | -10.47 |
| | | | | | | | | | | | 43 | -57 | -25 | FUSm(R) | -10.44 |
| | | | | | | | | | | | 32 | -43 | -8 | FUSm(R) | -10.41 |
| | | | | | | | | | | | 52 | -9 | -21 | MTGa(R) | -10.26 |
| | | | | | | | | | | | -42 | -82 | 13 | MTGp | -10.16 |
| | | | | | | | | | | | -25 | -49 | -18 | FUSm | -10.09 |
| 400-800 ms | -29 | -64 | -9 | FUSp | -10.82 | -30 | -54 | -26 | FUSm | -10.85 | 51 | -60 | -5 | ITG(R) | -13.64 |
| | -45 | -69 | -21 | FUSp | -10.66 | -58 | -44 | 14 | STGp | -10.59 | 38 | -59 | -40 | CBL1(R) | -12.58 |
| | 39 | -68 | -9 | FUSp(R) | -10.49 | -65 | -26 | -5 | MTGm | -10.58 | -31 | -73 | 29 | MOG/AG | -11.89 |
| | -52 | 8 | 25 | IFGop | -10.15 | -32 | -75 | 2 | FUSp | -10.47 | -56 | -40 | 7 | MTGp | -11.7 |
| | -42 | -69 | -11 | FUSp | -9.81 | -13 | -40 | -17 | FUSm | -10.37 | -54 | -36 | -18 | MTGp | -11.69 |
| | | | | | | 48 | -65 | -17 | FUSp(R) | -10.34 | 56 | -1 | -20 | MTGa(R) | -11.22 |
| | | | | | | -62 | -45 | -20 | ITGp | -10.25 | -49 | -49 | -22 | ITGp | -11.19 |
| | | | | | | -53 | -70 | -2 | MTGp | -10.22 | -46 | -64 | -1 | MTGp | -10.99 |
| | | | | | | -23 | -91 | 4 | FUSp | -9.9 | -27 | -50 | -19 | FUSp | -10.89 |
| | | | | | | -55 | -67 | -16 | ITGp | -9.86 | -39 | -79 | 22 | MOG/AG | -10.81 |
| | | | | | | -50 | -40 | 8 | MTGp | -9.86 | -51 | -58 | 16 | AG | -10.78 |
| | | | | | | -16 | -53 | -19 | FUSm | -9.83 | -3 | -43 | 40 | PCC | -10.71 |
| | | | | | | -41 | -71 | -4 | ITGp | -9.79 | -54 | -13 | -22 | MTGa | -10.6 |
| | | | | | | -48 | 11 | 28 | IFGop | -9.7 | 71 | -18 | 2 | STGm(R) | -10.52 |
| | | | | | | | | | | | -8 | -66 | 62 | PC | -10.3 |
| | | | | | | | | | | | 68 | -29 | 1 | STGp(R) | -10.19 |

| | | | | | | | | | | | | | | | |
|---|---|---|---|---|---|---|---|---|---|---|---|---|---|---|---|
| | | | | | | | | | | | -34 | -73 | -32 | CBL1 | -10.17 |
| | | | | | | | | | | | 41 | -15 | 7 | ROLop(R) | -10.04 |
| | | | | | | | | | | | -6 | 73 | 12 | FP | -9.91 |
| **800-1200 ms** | *No supra-threshold sources* | | | | | -58 | -36 | -6 | MTGm | -12.23 | -60 | -10 | 18 | ROLop | -10.42 |
| | | | | | | | | | | | -52 | -1 | 15 | ROLop | -10.35 |
| | | | | | | | | | | | -65 | -47 | 30 | SMG | -9.68 |
| | | | | | | | | | | | -55 | -7 | -16 | MTGa | -9.65 |
| **1200-1600 ms** | *No supra-threshold sources* | | | | | -54 | 3 | 19 | ROLop | -12.11 | -61 | -11 | 22 | ROLop | -11.34 |
| | | | | | | -60 | -8 | 16 | ROLop | -10.96 | -23 | -16 | -27 | PH | -11.26 |
| | | | | | | -7 | -47 | 15 | PCC | -10 | -65 | -14 | 11 | STGa | -10.08 |
| | | | | | | -65 | -24 | -4 | MTGm | -9.91 | | | | | |
| | | | | | | -67 | -32 | 11 | STGp | -9.9 | | | | | |
| **1600-2000 ms** | *No supra-threshold sources* | | | | | -43 | 2 | -12 | STGa | -10.17 | -43 | 2 | 12 | ROLop | -10.68 |
| | | | | | | -61 | -1 | 21 | ROLop | -10.13 | 51 | 19 | 15 | IFGop(R) | -10.36 |
| | | | | | | -60 | -6 | 27 | ROLop | -10.11 | | | | | |
| | | | | | | -68 | -29 | -11 | MTGm | -9.99 | | | | | |
| | | | | | | -45 | -16 | -16 | MTGm | -9.98 | | | | | |
| | | | | | | -35 | -5 | 15 | ROLop | -9.89 | | | | | |
| **2000-3000 ms** | *No supra-threshold sources* | | | | | -57 | 19 | 18 | IFGtr | -11.12 | *No supra-threshold sources* | | | | |
| | | | | | | -60 | -25 | 5 | STGm | -10.18 | | | | | |
| | | | | | | -35 | -5 | 15 | ROLop | -9.94 | | | | | |
| | | | | | | -63 | -29 | -6 | MTGm | -9.89 | | | | | |

**List of local maxima in classic frequency bands (Supplementary Table 2) and in the defined time windows (Supplementary Table 3).** For each of the three sentence parts (P1, P2, P3) and each frequency or time window, MNI coordinates (x, y, z) of local statistical maxima are given in mm, along with an approximate anatomical location and the corresponding *t*-value. Positive *t*-values indicate ERS whereas negative values represent ERD. A list of anatomical abbreviations is provided hereunder.

#### List of abbreviations

| Location | Suffix |
|---|---|
| **AG: angular gyrus**<br>**CAL: calcarine cortex**<br>**CBL1: cerebellum crus 1**<br>**FP: frontal pole**<br>**FUS: fusiform gyrus**<br>**MOG: middle occipital gyrus**<br>**PC: precuneus**<br>**PCC: posterior cingulate cortex**<br>**PH: parahippocampal gyrus**<br>**ROL: rolandic cortex**<br>**SOG: superior occipital gyrus** | **a: anterior aspect**<br>**m: middle aspect**<br>**p: posterior aspect**<br>**tr: pars triangularis**<br>**op: pars opercularis**<br>**or: pars orbitalis**<br><br>**<u>Side</u>**<br>All local maxima are located on the left side unless specified otherwise:<br>**(R): right-sided** |
